## Supplementary Figures for "Post-transcriptional control drives Aurora kinase A expression in human cancers"

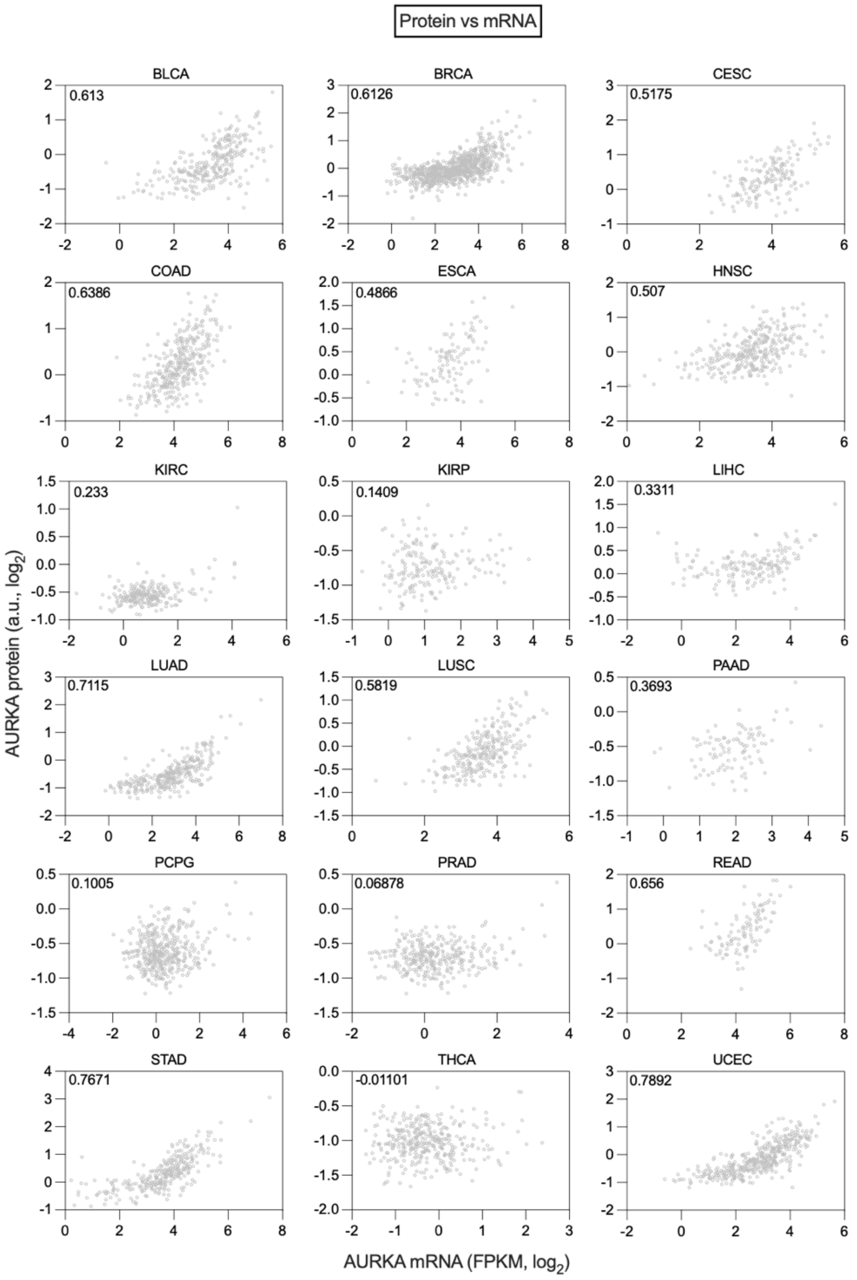


**Supplementary Figure 1.** Scatter plots displaying the correlation between AURKA mRNA and protein expression in 18 TCGA cancers. Data points represent individual tumour samples. Values of the Spearman's rank coefficient (r) shown for each cancer. p<0.001 for all r coefficients except PCPG and KIRP (p>0.05), PRAD and THCA (p=1). Note that the ranges of the x and y axes are different in each panel.

A

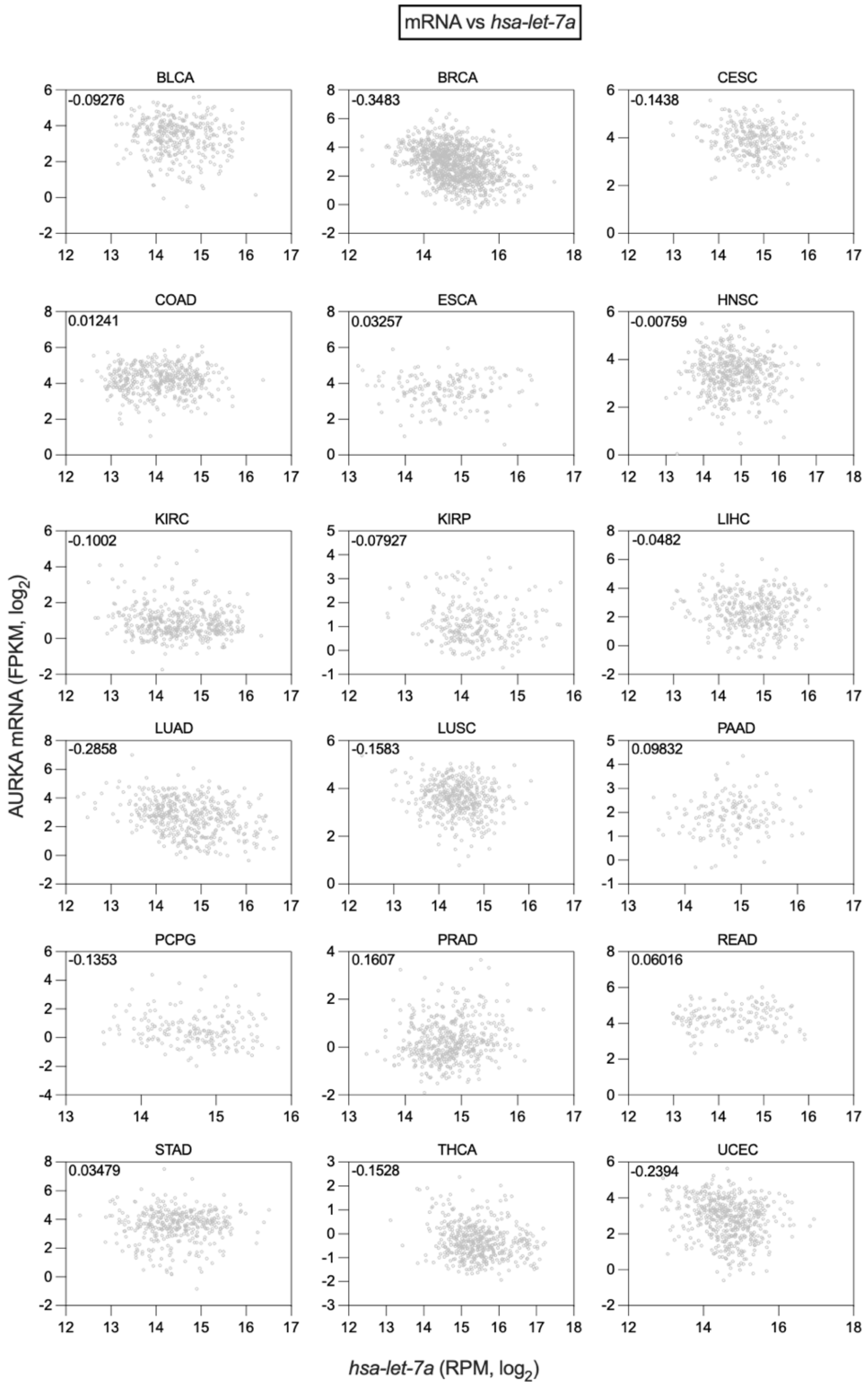
mRNA vs hsa-let-7a
BLCA
-0.09276
BRCA
-0.3483
CESC
-0.1438
COAD
0.01241
ESCA
0.03257
HNSC
-0.00759
KIRC
-0.1002
KIRP
-0.07927
LIHC
-0.0482
LUAD
-0.2858
LUSC
-0.1583
PAAD
0.09832
PCPG
-0.1353
PRAD
0.1607
READ
0.06016
STAD
0.03479
THCA
-0.1528
UCEC
-0.2394
AURKA mRNA (FPKM, log2)
hsa-let-7a (RPM, log2)

B

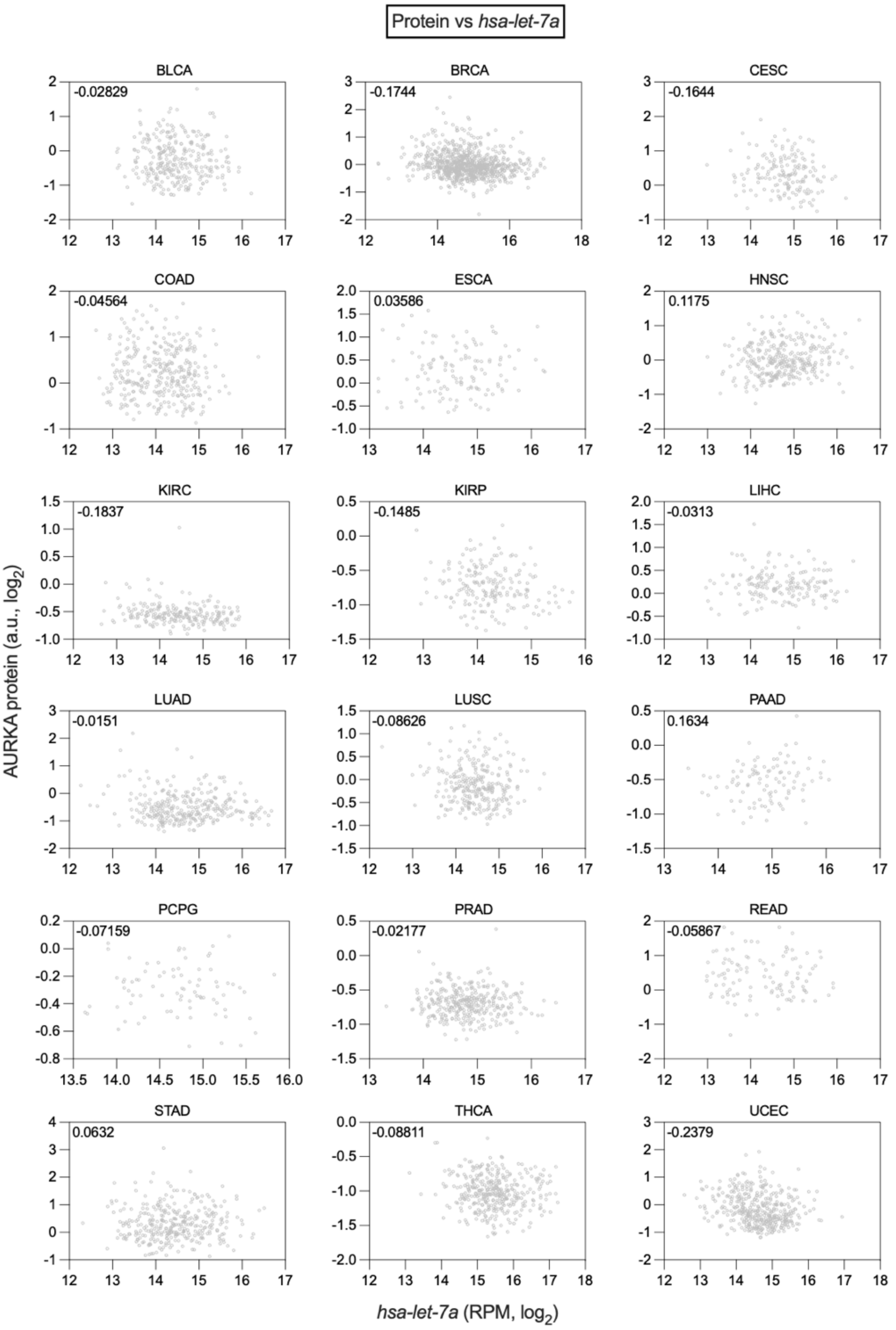


**Supplementary Figure 2. (A), (B)** Scatter plots displaying the correlation between expression of AURKA mRNA and *hsa-let-7a* (A) or AURKA protein and *hsa-let-7a* (B) in 18 TCGA cancers. Data points represent individual tumour samples. Values of the Spearman's rank coefficient (r) shown for each cancer. Note that the ranges of the x and y axes are different in each panel. **(A)** p<0.001 for r coefficients of BRCA, LUAD, PRAD, THCA, UCEC; p<0.05 for r coefficient of LUSC; p=1 for all other r coefficients except CESC (p=0.234) and KIRC (p=0.666). **(B)** p<0.001 for r coefficients of BRCA and UCEC; p>0.05 for r coefficients of CESC, HNSC, KIRC, KIRP; p=1 for all other r coefficients.

A

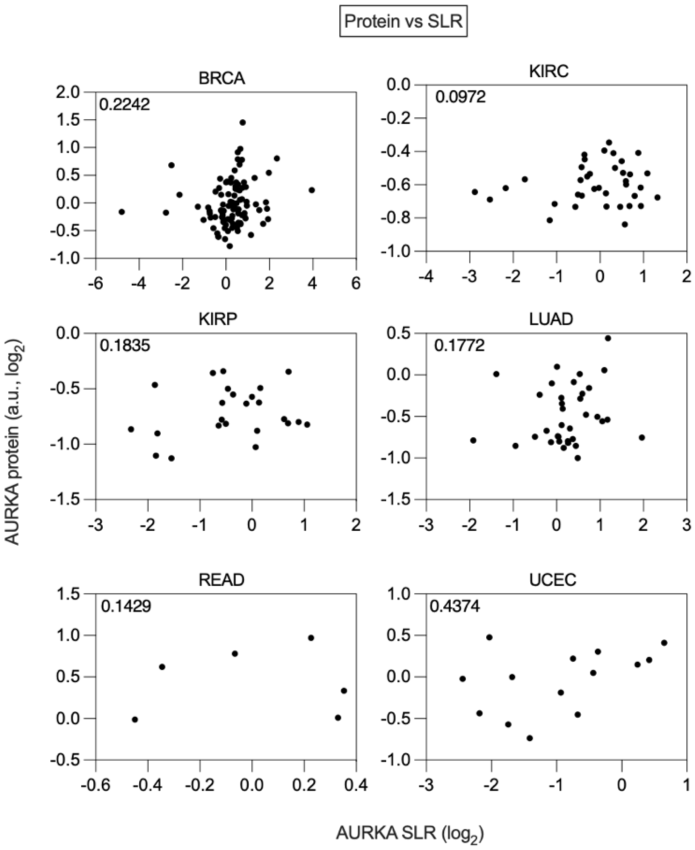


B

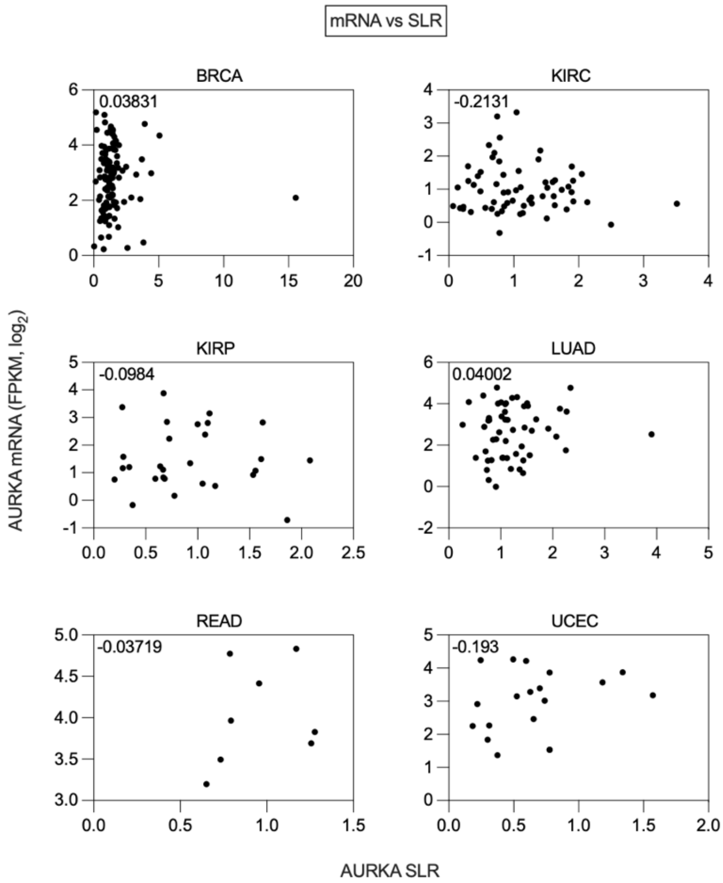


**Supplementary Figure 3. (A), (B)** Scatter plots displaying the correlation between AURKA SLR and expression of AURKA protein (A) or AURKA mRNA (B) in selected TCGA cancers. Data points represent individual tumour samples. Values of the Spearman's rank coefficient (r) shown for each cancer. Note that the ranges of the x and y axes are different in each panel.
